# A comparative study of influenza A M2 protein conformations in DOPC/DOPS liposomes and in native *E. coli* membranes

**DOI:** 10.1101/2024.01.08.574681

**Authors:** Griffin Sanders, Peter P. Borbat, Elka R. Georgieva

## Abstract

We compared the conformations of the transmembrane domain (TMD) of influenza A M2 (IAM2) protein reconstituted at pH 7.4 in DOPC/DOPS bilayers to those in isolated *E. coli* membranes, having preserved its native proteins and lipids. IAM2 is a single-pass transmembrane protein known to assemble into homo-tetrameric proton channel. To represent this channel, we made a construct containing the IAM2’s TMD region flanked by the juxtamembrane residues. The single cysteine substitute, L43C, of leucine located in the bilayer polar region was paramagnetically tagged with a methanethiosulfonate nitroxide label for the ESR (electron spin resonance) study. We compared the conformations of the spin-labeled IAM2 residing in DOPC/DOPS and native *E. coli* membranes using continuous-wave (CW) ESR and double electron-electron resonance (DEER) spectroscopy. The total protein-to-lipid molar ratio spanned the range from 1:230 to 1:10,400. The CW ESR spectra corresponded to a nearly rigid limit spin label dynamics in both environments. In all cases, the DEER data were reconstructed into the distance distributions showing well-resolved peaks at 1.68 nm and 2.37 nm. The peak distance ratio was 1.41±0.2 and the amplitude ratio was 2:1. This is what one expects from four nitroxide spin-labels located at the corners of a square, indicative of an axially symmetric tetramer.

Distance modeling of DEER data with molecular modeling software applied to the NMR molecular structures (PDB: 2L0J) confirmed the symmetry and closed state of the C-terminal exit pore of the IAM2 tetramer in agreement with the NMR model. Thus, we can conclude that IAM2 TMD has similar conformations in model and native *E. coli* membranes of comparable thickness and fluidity, notwithstanding the complexity of the *E. coli* membranes caused by their lipid diversity and the abundance of integral and peripheral membrane proteins.

## INTRODUCTION

Cellular membranes harbor active sites responsible for numerous physiological processes, and they provide the environments for proper folding of membrane proteins and assembly into functional units. *In vitro* studies typically utilize customized model membranes made of selected lipid types, which ensure known and relatively well-defined bilayer properties, e.g. hydrophobic thickness, charged vs. non-charged head groups, ordering, mechanical and dynamic properties, fluidity, and more. Consequently, customized lipid bilayers made of carefully chosen lipid compositions are routinely used to study membrane proteins.^1–3^ While lyso-lipid micelles and nanodiscs/lipodisqs are sometimes used to stabilize proteins for structural and in some cases functional assays,^1,4,5^ the studies of membrane proteins often require an ultimate test, that is one based on liposomes as the closest in form biological membrane mimetic.^1,6^ But how close are liposomes to native membrane as the medium of choice in studies of protein functional properties?

The lingering question is, how close protein conformations and dynamics observed in model bilayers are to what they may exhibit in native membrane under the conditions of lipid diversity, leaflet asymmetry, and protein crowding. Answering this question would require researching the different classes of membrane proteins in native membranes to learn and characterize the nature and extent of existing differences. Such highly desirable studies of membrane proteins (including homo-oligomeric proteins) can be carried out in native lipid environments,^7^ or as a minimum under approximating conditions, e.g., in native lipid mixtures that retain crowding and diversity of proteins populating the native membrane. A compelling example to that is the study of monoaminoxidases MAO-A and MAO-B dimers stability in outer mitochondrial membrane fragments and lipid mimetics.^8^ Therefore, an interesting class of proteins that can be studied vis-à-vis their response to environment are transmembrane homo-oligomeric proteins which attain specific function upon assembly and can have properties that can be modified by the specificity of biomembrane environment.

Here, we subjected to test the assembling and conformation of the transmembrane domain (TMD) of the influenza A M2 proton channel (IM2)^9–12^ residing in model membranes of DOPC/DOPS (1,2-dioleoyl-sn-glycero-3-phosphocholine/1,2-dioleoyl-sn-glycero-3-phospho-L-serine) vs. isolated *E. coli* membranes with most of native proteins preserved. It would be hard to retain all of the peripheral membrane proteins but they do not constitute a major protein fraction associated with *E. coli* membranes.^13^ IM2 functions as a tetrameric proton-conducting channel aiding virus adaptation and proliferation.^11,12^ Previously, we studied the assembly of IM2 TMD in liposomes made of DOPC/POPS (1-palmitoyl-2-oleoyl-sn-glycero-3-phospho-L-serine) using spin labeling and pulse electron spin resonance (ESR) spectroscopy, namely DEER (double electron-electron resonance).^14,15^ Based on the concentration dependence of modulation depth (Δ) of the DEER signal recorded over two orders of magnitude range of protein-to-lipid molar ratios (P/L),^14,16,17^ we concluded that the protein tetramer assembled via a tandem mechanism, which includes initial monomer-to-dimer self-association, and at the upper range of P/L IM2 tetramer was a dominant form.^14^ In that work we used spin-labeling of L46C peptide containing the IM2 TMD flanked by the N- and C-terminus juxtamembrane residues.^14,15^ The effect of lipids and pH on assembly was observed, but the structural implications remained elusive.

Here, we used again spin-labeled IM2 TMD and studied it by continuous-wave (CW) ESR and DEER methods.^18,19^ The IAM2 peptide composition and length were close to those used previously,^14,15^ but the spin-labeled residue was L43C rather than L46C used previously. The L43C residue is right at the C-terminus of the IM2 TMD helix located in the headgroup region of the membrane bilayer (Figure 1), so that when studied by ESR it informs on the conformation and oligomeric state of this protein region, which is believed to be a better reporter of protein structure and dynamics. Remarkably, the ESR data for spin-labeled IM2 TMD in DOPC/DOPS liposomes and in native *E. coli* membranes were almost identical, which suggests similar conformations of IM2 in these environments and thus excellent capacity of selected liposomes to represent *E. coli* lipid environment in the case of IM2, at least at the first glance. The CW ESR spectra for all samples were rather broad, suggesting highly restricted motional dynamics of the spin label and the protein region itself in the proximity to the residue L43C.^20^ The conformational restraints resulted in clearly oscillating DEER signals originating from the dipolar interaction between spin labels at L43C sites; the DEER signals were reconstructed into well-defined bimodal distance distributions corresponding in their major part to the side and diagonal inter-spin distances of spin labels in symmetric IM2 tetramer. The dominant two narrow peaks in distance distributions were at 1.68 nm and 2.37 nm. The peak distance ratio was 1.41±0.2 and the amplitude ratio was 2:1. This is exactly what one expects from four nitroxide spin-labels situated at the corners of a square, supporting an axially symmetric tetramer. Consistent features at ∼2.0 nm and minor peak 2.8 nm could also be noticed, but finding their exact origin was not a goal of this work.

**Figure 1.**
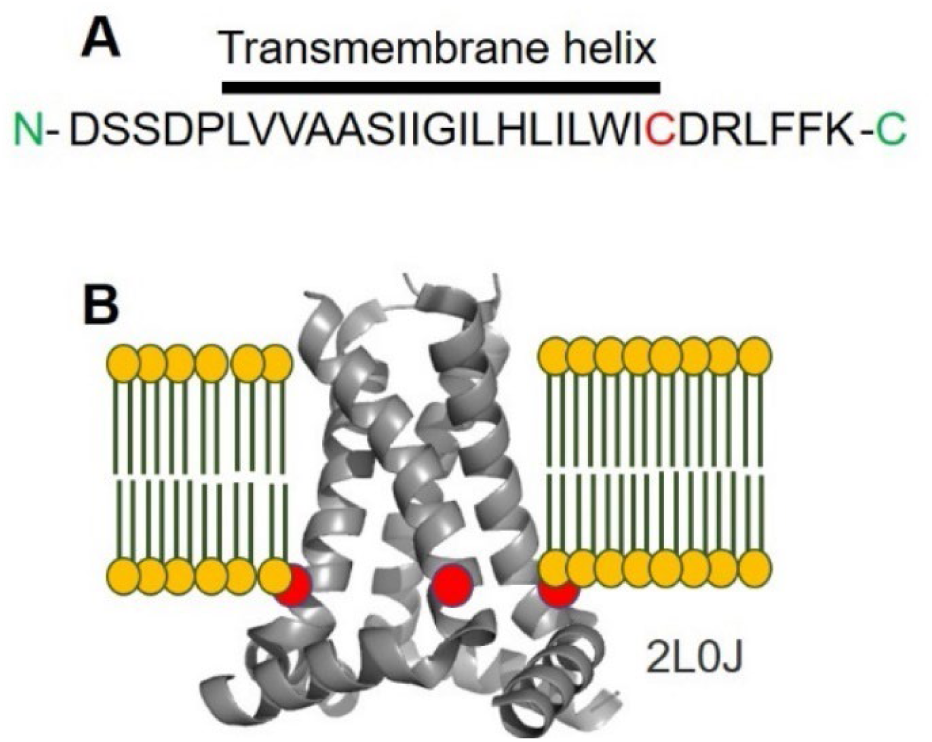
IM2 amino acid sequence and structure: (A) The amino acid sequence of the M2 construct used in this study. The transmembrane domain (TMD) is indicated with overline, and the cysteine residue (L43C, amino acid numbering in full-length M2) is in red color. (B) The solid-state NMR structure of M2 TMD and C-terminal soluble helix (PDB: 2L0J) is shown as a cartoon within the bilayer. The location of the L43 residue is indicated with a red circle.

Thus, in this work, we present a simple economical protocol for reconstitution of spin-labeled single-pass transmembrane protein IM2 into membranes made of native lipids and under conditions of protein crowding. We successfully assessed the IM2 tetramer in these environments by using site-directed spin labeling (SDSL) and ESR spectroscopy. The closeness of the data for IM2 in DOPC/DOPS and *E. coli* membranes suggests that the DOPC/DOPS bilayer provides an adequate environment to assess IM2’s structure and possibly its function.

We also elaborated the study by comparing the data obtained on *E. coli* inner membrane (IM) extracts to the total mix that also contained outer membrane (OM). Furthermore, in addition to previous studies,^14,20–25^ our results unambiguously demonstrate the utility of ESR to assess protein conformations and of pulse DEER spectroscopy in providing accurate distances within an oligomeric protein under diverse membrane conditions. Our study validates the utility of pulse DEER spectroscopy in providing accurate distances within an oligomeric protein under diverse membrane conditions.

## MATERIAL AND METHODS

### IM2 TMD synthesis, solubilization and spin labeling

The peptide corresponding to the IAM2 TMD was commercially synthesized (GenScript, Inc.) and received as a powder. Then, it was solubilized in a buffer containing 20 mM Tris/HCl pH 7.4 150 mM NaCl, 5 mM DTT (Dithiothreitol) and 10 mM of the detergent β-DDM (n-Dodecyl-β-D-maltoside) and incubated for 1 h at room temperature (RT) under constant agitation. The solubilized and refolded peptide was dialyzed against the same buffer but with 1 mM β-DDM and no DTT to decrease detergent concentration and remove DTT. Thereafter, the 100-120 µM IM2 TMD peptide solution was mixed with the cysteine-specific methanothyosulphonate spin label (MTSL) at a 1:30 molar ratio and the reaction between the L43C residue and MTSL was conducted at RT for 3-4 h and then at 4 °C overnight, as previously described.^14,15^ The unreacted spin label was removed by dialysis against the buffer containing 20 mM Tris/HCl pH 7.4, 150 mM NaCl and 1 mM β-DDM. The final concentration of spin-labeled IM2 TMD was 78 µM or 107 µM, estimated by the absorbance at 280 nm,^14^ and these stock solutions were further used for IM2 TMD reconstitution in lipid membranes.

### Production and isolation of native *E. coli* membranes

*E. coli* membranes were isolated after growing *E. coli* BL21(DE3) cells transfected with a plasmid pET15a having cloned the gene encoding the HTLV-1 p13^II^ protein.^26^ No p13^II^ overexpression was initiated as the plasmid was one at hand serving only to provide the *E. coli* cells with ampicillin (Amp) resistance. The cells were grown first at 37 °C for 3 h and overnight at 14 °C, harvested the next day and broken open. The membranes were isolated with ultracentrifugation.^21^ These membranes contained the native *E. coli* lipids and membrane proteins. *E. coli*’s inner membranes (IM) used in control DEER experiment, were separated from outer membranes using a modified gradient ultracentrifugation protocol by Shu and Mi^27^ as described in the Supplement. These membranes (native *E. coli* lipids and membrane proteins) were used for reconstitution of spin-labeled IM2 TMD.

### Preparation of DOPC/DOPS lipid mixture and *E. coli* membranes

DOPC and DOPS, in chloroform and chloroform/methanol/H_2_O, respectively, were purchased from Avanti Polar Lipids, Inc. Then, aliquots were taken from these lipid solutions and mixed to produce the molar ratio of 70% DOPC and 30% DOPS. The solvent was evaporated under a stream of nitrogen gas until no solvent traces were visible and then was flashed with nitrogen gas for additional 2 h to remove the residual organic solvent. A buffer of 20 mM Tris/HCl pH 7.4, 150 mM NaCl and 1 mM EDTA was added to obtain a final total lipid concentration of 30 mM. The dried lipids were incubated with the buffer for 1 h at 4 °C to complete lipid hydration, and the mixture was vortexed to obtain homogeneous lipid distribution in the solution volume. This lipid mixture was used as a stock solution to prepare the IM2 TMD liposome samples.

### Reconstitution of the spin-labeled IM2 TMD in DOPC/DOPS liposomes and native *E. coli* membranes

The IM2 TMD reconstitution was conducted as follows: First, wet *E. coli* membranes were weighted and solubilized in a buffer of 20 mM Tris/HCl pH 7.5, 150 mM NaCl and 1 mM EDTA. In parallel, aliquots of the 30 mM DOPC/DOPS stock solution were taken. The lipid membranes were solubilized using a 10% solution of Triton X-100 in water by adding aliquots of the Triton X-100 solution to the membrane solutions until they become visibly transparent. Thereafter, spin-labeled IM2 TMD protein was added to each lipid sample to obtain the desired P/L. The mixtures were incubated for 1 h at RT and then BioBeads (BioRad) were added to each sample and incubated for 2 more hours under constant agitation to remove the detergents. The BioBeads were changed, and the samples were transferred to 4 °C to stay for 2 h. The procedure was then repeated 3 times for a total of 16 h. The next day, the BioBeads were removed and the resulting IM2 TMD-liposome samples were concentrated and used for ESR measurements.

The incorporation of IM2 into *E. coli* membranes was conducted in similar way. After membrane treatment with 10% solution of Triton X-100, the spin-labeled IM2 TMD was added to the processed membranes. Several samples in a range of P/L’s were prepared. Sample numbering, compositions, and estimates of P/L ratios for *E. coli* membranes are compiled in the Supplementary Table 1 alongside with the IM2 TMD samples in DOPC/DOPS.

### Pulse EPR (DEER) at 17.3 GHz on spin-labeled IM2 TMD in lipid membranes, inter-spin label distance reconstructions and modeling of the distances based on existing IM2 structures

The pulse ESR (DEER) measurements were conducted at ACERT, Cornell University. The measurements were carried out at 60 K using a home-built Ku-band pulse ESR spectrometer operating at MW frequency band around 17.3 GHz.^28,29^ A DEER experiment setup^22^ used for electron spin-echo detection a π/2-π-π pulse sequence with 32-ns π-pulse applied at the low-field edge of the nitroxide spectrum. A 16-ns π-pulse at a 70 MHz lower frequency pumped at the spectrum central peak. A 32-step phase cycle^30^ was applied for suppressing unwanted signal contributions to the refocused echo and some other instrumental artifacts, leaving only three unwanted minor dipolar pathways produced by the 4-pulse DEER sequence. Such contributions caused by dipolar coupling are phase-independent and cannot be removed completely.^31^ Nuclear electron spin-echo envelope modulation (ESEEM) caused by the matrix protons was suppressed by summing up the data traces recorded in four sequential measurements in which initial pulse separations and the start time of detection were advanced in the subsequent measurement by 9.5 ns that is by a quarter period of the 26.2-MHz nuclear Zeeman frequency of protons at 615 mT corresponding to 17.3 GHz working frequency.^32^

The recorded raw DEER signals (Supplementary Figures 1-3) were background corrected, the end points of the records containing visible distortions caused by unwanted dipolar pathways and instrumentation artifacts were omitted, and the distances were reconstructed using Tikhonov regularization software or SVD, as needed.^33,34^ The reconstructions were stable and similar with both methods.

Molecular modeling software (MMM)^35^ was used to generate spin label’s (MTSL’s) rotamer libraries and to predict distances and distance distributions based on existing NMR and X-ray structures of IM2 TMD. The predictions were carried out for conditions at cryogenic temperatures.

It would not be unusual to encounter some level of spin-label reduction at low protein concentrations which could reduce DEER modulation depth, at low P/L ratios. To estimate if this was significant, primary echo amplitudes were measured for all samples. The assessment of spin label concentration vs. estimated protein concentrations was made using DEER and primary echo data for samples 1a-5a by first measuring primary echo (PE) amplitudes using 16 ns and 32 ns pulses separated by 250 ns at low-field edge of the nitroxide spectrum and 100 Hz pulse repetition rate, scaling, and introducing known corrections. The amplitudes were scaled using P/L ratios and corrected to the loss of primary echo amplitude to spin-echo dephasing caused by dipolar interaction by applying the factor of (1-*p*)^-1^. Here *p* is loss of primary echo amplitude and is proportional to the modulation depth Δ in DEER. The dipolar dephasing time was nearly complete within the 250 ns pulse separation in primary echo. The results are compiled in Supplementary Table 2 and indicated that the contribution from spin label loss is not a significant contribution to Δ versus P/L profile.

### CW EPR measurements on spin-labeled IM2 TMD in lipid membranes

CW ESR experiments were carried out using an X-band Bruker E-500 spectrometer. The spectra were recorded at 10±1 °C using magnetic field modulation amplitude of 0.2 mT and 2 to 2.5 mW microwave power.

## RESULTS

### The CW ESR spectra of spin-labeled L43C residues of IM2 TMD in DOPC/DOPS and native *E. coli* membranes are similar

For all studied samples of IM2 TMD spin-labeled at residue L43C, we recorded close in shape and broad CW ESR spectra as shown in Figure 2. Spin label is highly immobilized in spin label/protein region in the proximity to the membrane surface.^23,24,36^ These spectra were also much broader than the previously recorded CW ESR spectra for residue L46C in liposomes,^15^ reflecting the differences in the dynamics of these residues with L43C being highly restrained compared to L46C case. Interestingly, the outer splitting in CW ESR spectrum for residue L43C is smaller in DOPC/DOPS vs. *E. coli* membranes (Figure 2), possibly due to less polar environment of the spin label in the model liposomes.

**Figure 2.**
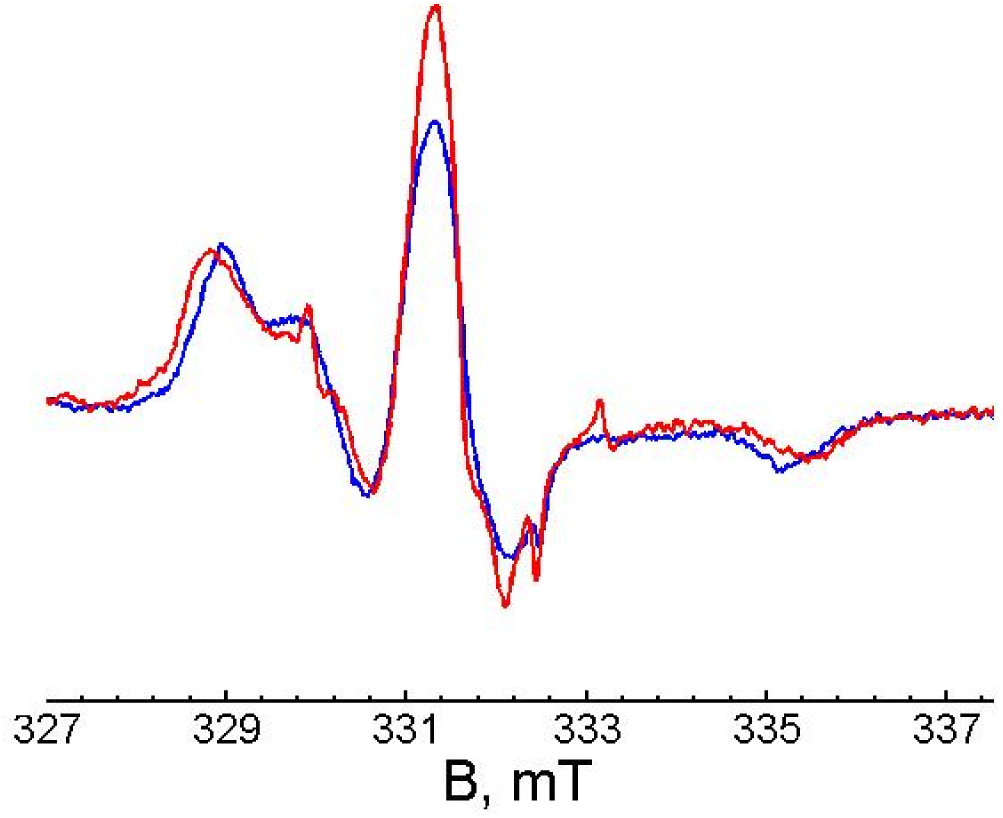
CW ESR spectra of IM2 TMD spin-labeled at residues L43C. IM2 TMD in DOPC/DOPS liposomes (red); IM2 TMD in native *E. coli* membranes (blue). The low-intensity narrow lines are caused by a very small fraction of unreacted free spin label, which does not otherwise affect the spectrum of protein-attached spin label. The observed CW ESR spectra correspond to a very slow-motion range nearing rigid limit of the spin label and thus underlying protein region dynamics. Note that in liposomes the central component is broadened most likely by dipolar coupling between spin-labels due to a prevalence of tetramers with ≈1.7 nm distances between proximal labels. (A narrow feature at 332.6 mT is from E′ centers in fused silica sample tubes.)

### The time-domain DEER data and reconstructed inter-spin label distances suggest similar IM2 TMD tetramer conformation in DOPC/DOPS and native *E. coli* membranes

The DEER data recorded for all the samples of IM2 TMD with spin-labeled residue L43C in DOPC/DOPS and *E. coli* membranes were remarkably similar and clearly contributed by two dominant inter-spin distances (Figure 3A). Furthermore, due to the location of the spin-labeled residue L43C at the solvent/membrane interface, which restricts the conformational heterogeneity of this protein region and possibly the number of populated MTSL rotamers,^37^ all DEER data are of well-defined oscillatory type. This allows reconstruction (Figure 3B, C) of narrowly-bounded distances with main peaks at *R*_1_=1.68 nm and *R*_2_=2.37 nm (*R*_2_ ≍ 1.41 *R*_1_), with distances in a ratio of 2^1/2^ corresponding to the diagonal and side of a square as expected for a symmetric tetramer.^14,38^ This is well in line with the previous finding for IM2 TMD spin-labeled at residue L46C.^14^ In this earlier study,^14^ however, the used residue L46C is at the surface of the lipid bilayer and is more exposed to the solvent, permitting greater conformational heterogeneity of the corresponding protein region and higher spin label mobility; yielding less resolved distances contributed by the IM2 tetramer. Therefore, the current data for residue L43C report much more precisely on the arrangement of IM2 transmembrane helices within the IM2 tetramer and have potential of reporting on other possible oligomeric constructs (e.g. dimers) or specific effects of lipid and pH. For all samples a slightly varying in intensity peak at a distance of about 2.8 nm could be noticed as well (Figure 3B and C). It is possibly contributed by IM2 TMD dimers, because a dimer was found to be an intermediate stage in the process of IM2 tetramer assembly.^14^ This could also correspond to a more open conformation/state of the channel reported by X-ray structures and possibly one populated at low pH, as we have a moderate confidence evidence for that at a low-pH *E. coli* inner membrane (IM) sample as shown in the **Supplementary figure 5**.

**Figure 3.**
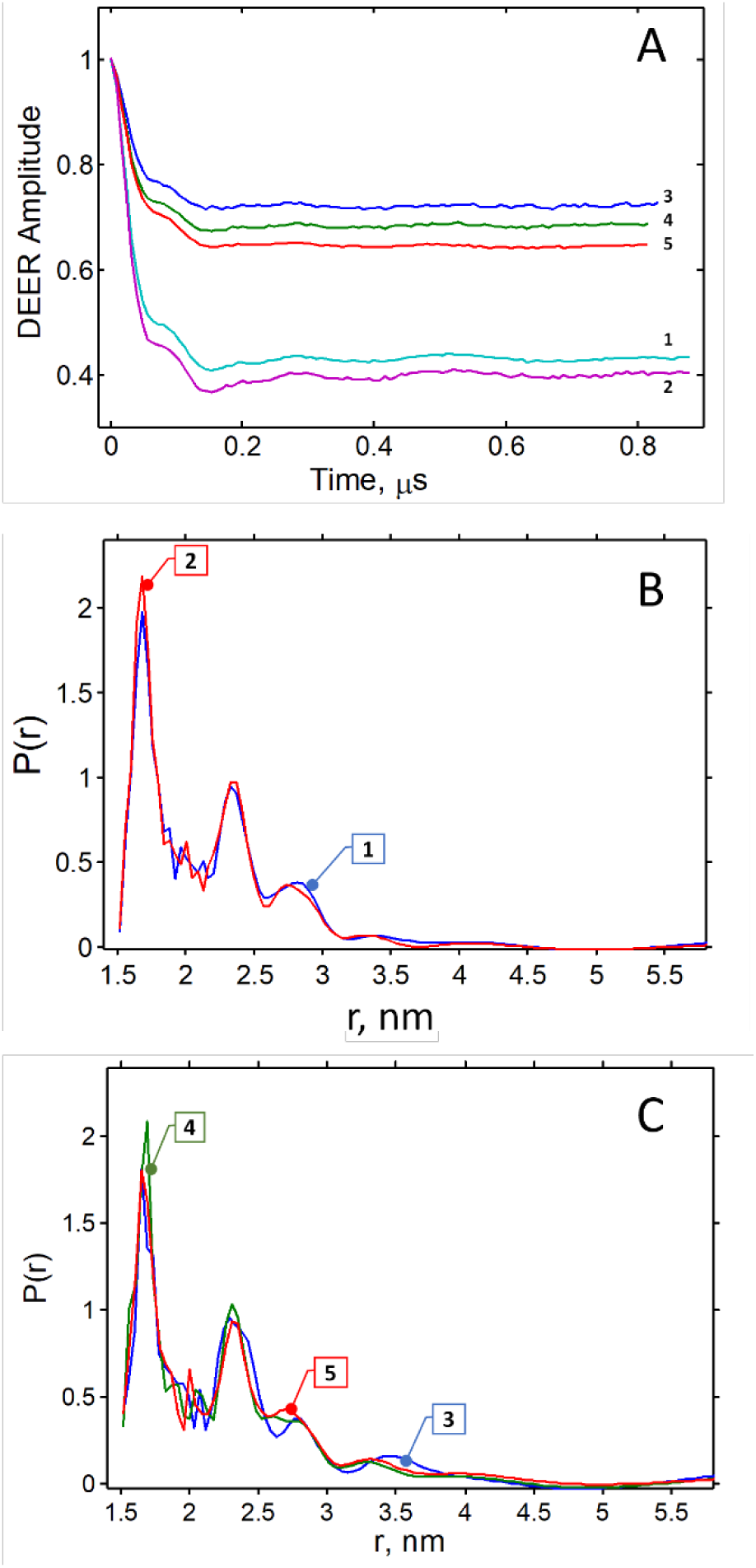
DEER data for the IM2 TMD spin-labeled at L43C residues in DOPC/DOPS and native *E. coli* membranes. (A) Baseline-corrected time-domain DEER data for all samples; The numbers are that denoting samples in the **Supplementary Table 1** where the PLR’s and sample descriptions are also given. (B) Reconstructed DEER distances for IM2 TMD in DOPC/DOPS (C) Reconstructed DEER distances for IM2 TMD in the native *E. coli* membranes. The numbers in (B) and (C) correspond to the sample numbers.

Two samples of IM2 TMD in DOPC/DOPS with P/L’s of 1:480 and 1:230—Sample 1 and Sample 2, respectively—were measured. We found that the DEER’s modulation amplitudes Δ in Figure 3A for Sample 2 is about 0.6, which is close to expected 0.725 in the ideal case of fully-assembled tetramer with each monomer containing spin label. The Δ can be estimated as Δ(*p*, *N*) = [1 – (1– *p*) *^N^*^−1^],^14,16,17^ where *N* is the number of spin-labeled monomers (assuming 100% spin labeling efficiency) and using well-known depth *p* of 0.35 for the 16 ns pump pulse for the DEER on a standard biradical using the settings in this work. Of course, Δ for Sample 1 with a P/L of 1:480 is less than in the ideal case (of a fully labeled 100% tetramer) caused by a fraction of IM2 TMD dimers at this P/L^14^ and less than 100% spin labeling further reducing Δ. At the same time, the DEER Δ’s for samples of IM2 TMD in *E. coli*’s membranes (Samples 3, 4 and 5) were all smaller than those for the samples in DOPC/DOPS liposomes (Figure 3A vs. 3B), however the P/L’s were estimated as greater than those for IM2 TMD in liposomes. Furthermore, in an additional series of DEER measurements on samples of IM2 TMD in *E. coli* membranes having P/L’s ranging from 1:400 to 1:10,400 (Samples 1a-5a in Supplementary Table 1), the DEER Δ drops significantly with the decrease of P/L ratio (Supplementary Figure 2). Additionally, the population of the distance at 2.8 nm increased upon decreasing the P (IM2 TMD)/L and becomes dominating for Sample 5a with P/L of about 1:10,400. (It should be noted that the lower SNR at high P/L ratios could negatively affect the quality of reconstructed distances distributions). Thus, the DEER results from the current study are suggests that IM2 TMD tetramer in *E. coli* membranes assembles via a dimer intermediate in a qualitative agreement with what was found previously for this protein in model bilayers.^14^ However, the observed decrease in Δ did not have significant effect on the shape (form-factor) of the DEER signal and the reconstructed inter-spin distance distributions were very close to those for IM2 TMD in model membranes, particularly for P/L’s below 1:600 (Supplementary Figure 2). Other factors, which could lead to the decreased DEER Δ might be the partial spin-label reduction or detachment from the protein at the conditions encountered during the reconstitution of the spin-labeled IM2 TMD in *E. coli* native membranes, given that the native still functional proteins and co-factors and possibly other active native compounds were present. To address this concern, the primary echo amplitude analysis was carried out on a series of similarly prepared samples of IM2 TMD in *E. coli* membranes with P/L ratios from 400 to 10,400 obtained using the same stocks (Samples 1a to 5a as described in the Supplementary Information). We found that primary echo amplitudes (which depend on concentration of electron spins in the sample) for each of these samples were consistent within rmsd of 7% with the estimated protein concentrations (Supplementary Tables 1 and 2). Also, negligible fractions of free spin label in the samples were detected by CW ESR. These results suggest that there is no significant loss of spin label under our experimental conditions for IM2 TMD in *E. coli* membranes. One should also keep in mind that the high content of native membrane proteins could affects protein lateral diffusion and binding affinity. This might also cause a shift in the equilibrium of IM2 TMD oligomer due to slower diffusion of IM2 monomers caused by boundary lipids and to native protein-created kinetic traps in the native membrane.^39^

Further, we reconstituted the spin-labeled IM2 TMD in isolated *E. coli* inner membrane (IM) to serve as a control, because we did not spend effort to separate the fraction of remaining outer membrane (OM) in most of the preparations. The outer leaflet of BL21 OM is mostly formed by LPS^40^ and is not very likely to have IM2 present there. Internal leaflet of OM has similar lipidome to IM^40^ and the OM may contain patches of lipid bilayer where IM2 could be inserted, this would be similar to IM but most likely is insignificant contribution. It is also possible that much of OM was shed along with the peptidoglycan sacculus. So, while reconstitution of IM2 could in principle take place in OM, our DEER results for spin-labeled IM2 TMD in the whole membrane extract were not any different from those obtained for protein in just IM (Supplementary Figure 4), suggesting that the OM most likely did not play a role in our sample preparation method.

### Molecular modeling of distances in existing IM2 tetramer structures vs. DEER experimental distances for spin-labeled residues L43C suggests the lipid environment plays a role in stabilizing protein’s conformation

To gain more insights into the structure of IM2 TMD, we used the molecular modeling software MMM^35^ to generate an MTSL rotamer library for the residue L43C based on existing IM2 tetramer structures. The best agreement between experimental and predicted distance distributions was achieved for the solid state NMR (ssNMR) IM2 structure at pH 7.5 in aligned 1,2-dioleoyl-sn-glycero-3-phosphatidylcholine:1,2-dioleoyl-sn-glycero-3-phosphoethanolamine bilayers (DOPC/DOPE) (Figure 4).^41^ Still, despite being close, the predicted and experimentally obtained distance distributions are not quite the same, because the MMM analysis predicted significantly broader distance distributions than those reported by DEER and possibly deemphasized dominant rotamers. Although the prediction and experiment produced bimodal distance distributions, the predicted maxima at about 2 nm and 3 nm (Figure 4 B) are shifted to longer distances compared to the experimental peak maxima at 1.68 nm and 2.37 nm (Figure 3 B and C). One possibility is that the MMM may have predicted a wider range of MTSL rotamers when attached to residue L43C (Figure 4) even at cryogenic temperature, but these rotamers become much more selectively populated under our experimental conditions, due to the presence of lipids not considered by MMM, thus producing narrower experimental distance distributions. Most likely the χ1-χ3 dihedral angles of MTSL side-chain are restrained, leaving some freedom to χ4/χ5 to present “tether-in-a-cone” case.^42^ DEER supports conformational ordering and tight packing of this protein region, as inferred from CW EPR (Figure 2).

**Figure 4.**
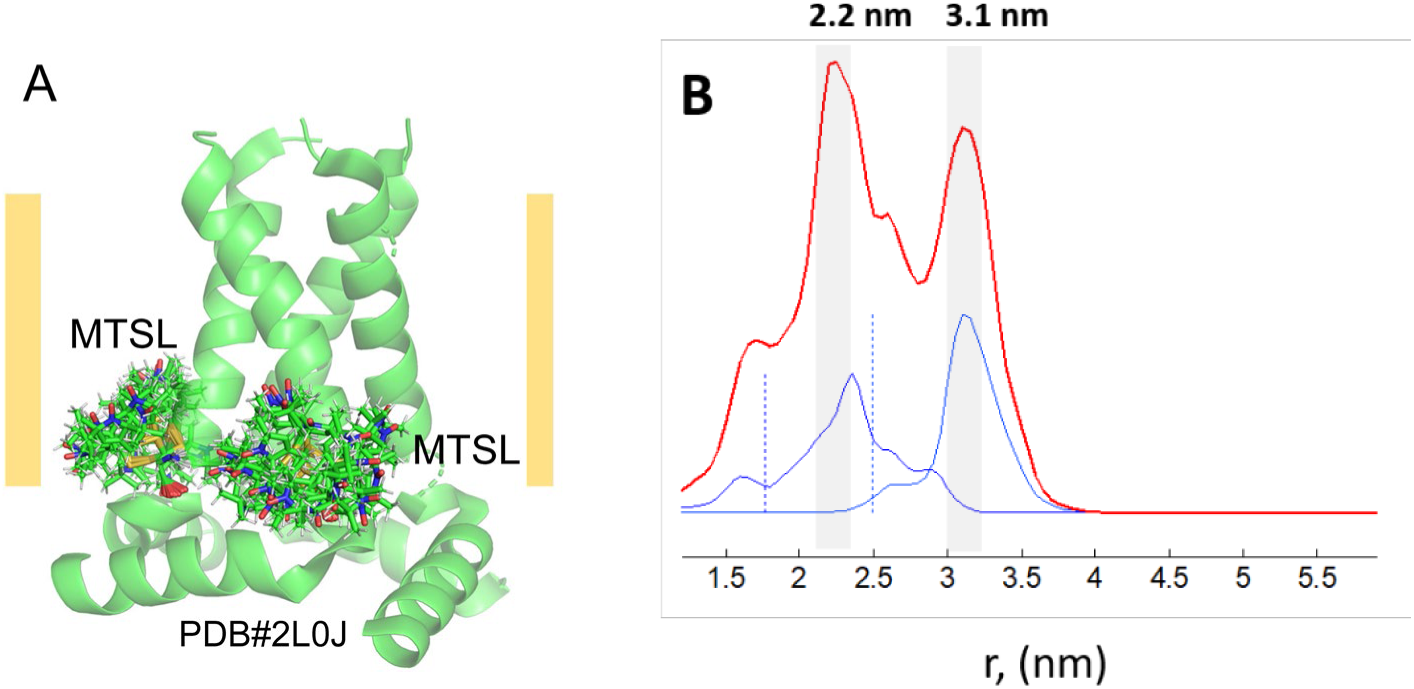
MMM analysis of distance distributions between spin-labeled residue L43C within IM2 tetramer. (A) The ssNMR structure (PDB: 2L0J) was used as a template to generate MTSL rotamer libraries and predict the inter spin-label distances. The MTSL rotamers for only two IM2 protomers are shown. (B) Based on the MMM analysis and predicted MTSL rotamers, bimodal distance distribution was obtained with maxima at about 2.2 nm and 3.1 nm.

We also used available IM2 X-ray structures to predict the distances and distance distributions based on MMM-generated MTSL rotamer libraries, but most of these distributions were significantly outside the ranges of our experimental DEER distances at physiological pH. Thus, for IM2 TMD’s crystal structure in lipidic cubic phase (LCP) at a high pH,^43^ MMM produced distances with maxima at up to 4 nm, which is much longer than the main peaks of our DEER distances within IM2 TMD tetramer (Supplementary Figures 2 and 3), suggesting different protein conformations in lipid bilayers vs. LCP. The reconstructed distances in Figure 3 contain a minor contribution of distances in the range of observed in cubic phase, but further studies would be needed to clarify if they really originate from alternative IM2 TMD alternative more open conformation.

The only observation in this work in support of such long distances was obtained for *E. coli* membrane sample adjusted to pH 5.4 by substitution of the Tris/HCl buffer with MES. For this samples, in addition to the DEER distance for IM2 TMD in closed conformation, additional pair of longer distances at 2,8/3.9 nm was observed (Supplementary Figure 5), which might represent the IM2 TMD open C-terminal pore, in agreement with previous results,^14,15^ and might also represent closer the X-ray structure. However, this would require further more detailed investigations and is beyond the scope of the present work.

## DISCUSSION

*In vitro* studies contribute a wealth of information on the structure and functions of integral membrane proteins. Typically, these studies are conducted on proteins stabilized by detergents, LPCs or residing in model lipid bilayers of controlled composition including special cases of bicelles and nanodiscs.^1,23,44,45^ These environments to a varying degree are acceptable substitutes for native membranes in that they stabilize membrane proteins for structural studies carried by NMR, ESR, X-ray crystallography, or electron microscopy. Liposomes are particular membrane mimetics, which provide some or most of the following properties: hydrophobic environment, true bilayer morphology, selection of lipid hydrocarbon chains length and bond saturation, and the type of lipid headgroups. However, their makeup is usually substantially simpler than the composition of native membranes and many synthetic lipids may prove costly for routine work. Also, these studies are normally conducted on protein of interest reconstituted in detergent or lipid with no other proteins being present. But native membrane’s morphology is determined to a large extent by the diverse lipid composition and proteins either traversing or linked to the membrane. Therefore, there is not well explored subject of how these membranes, comprising native lipids and crowded proteins, may affect integral membrane protein structure, dynamics, and subunit assembly. It is reasonable to expect, that the lateral movement may be affected and could even result in confinement due to percolation or other mobility obstructions; translational and rotational diffusion may slow down due to the presence of boundary lipids^46,47^ while interactions with transmembrane domains and loops of other crowded integral proteins may lead to kinetic trapping and non-Brownian diffusion.^48^ This may significantly affect oligomer assembly. Also, it is not fully understood how to select model membranes for in vitro studies to assess the structures and higher energy states that may be populated and be active under native conditions.

In this work, we compared the conformations of IM2 TMD in model DOPC/DOPS lipid bilayers and in native *E. coli* membranes. Although IM2 is expressed and functions in mammalian host’s membranes, studying this protein in *E. coli* membranes is worthwhile, as it was previously proposed that under native conditions the protein avoids cholesterol-rich domains,^49^ and it was also found that the influenza A envelope contains high amounts of PE lipid,^50^ which is also the major lipid in *E. coli* membranes.^51^ PE is also one of the main structural components of mammalian cell plasma and endomembranes.^52^ Furthermore, we included PS lipid because it is a component of influenza A envelope,^50^ and PS and PC (to a greater extent than PS) are components of host’s membranes as well.^52^ It was also suggested that positively charged amino acids in the C-terminal region of IM2, e.g., R45 and K49 (Figure 1), which are present in the studied here IM2 TMD construct, could interact with negatively charged lipid headgroups,^53^ notwithstanding this was later mostly deemphasized.^54^ The phosphatidyl glycerol (PG) lipid present in *E. coli*’s membrane^51^ makes these negative charges available as a substitute for PS in mammalian membranes, where IM2 resides. Therefore, in terms of lipid composition, all the above considerations are supportive of physiological relevance of the lipid membranes used here.

Previously, different constructs of IM2 containing just the TMD or TMD and the following amphipathic helix (Figure 1) in different lipid environments were studied mostly by NMR^41,55,56^ and X-ray crystallography.^43^ In these studies different lipid compositions were used, namely, PC lipid,^55,57^ PC/PG mixture,^41^ or compositions that are close to the viral envelope.^55,58^ One general observation in these earlier studies was that the IM2 tetramer stability and conformation depends on the lipid bilayer hydrophobic thickness, which we also observed in our past PDS study.^15^ Thus, it was found by NMR that the IM2 TM helix tilt varies for protein in lipids with a shorter vs. longer hydrocarbon chain, e.g., the tilt angle with respect to the membrane director for IM2 in DMPC (1,2-dimyristoyl-sn-glycero-3-phosphocholine) was somewhat greater than it was in DOPC bilayers.^57^ In our previous studies, we found that the IM2 tetramer assembles more efficiently in lipid bilayers made of DOPC/POPS than in DLPC/DLPS (1,2-dilauroyl-sn-glycero-3-phosphocholine/1,2-dilauroyl-sn-glycero-3-phospho-L-serine) lipids.^15^ These results emphasize the role of hydrophobic matching between IM2 TM helix and the membrane bilayer thickness. In terms of hydrophobic matching, DOPC/DOPS provides sufficient thickness to fully accommodate the IM2 TM helix of 20 amino acids.^59^ This possibly explains the almost identical CW ESR and DEER results for the spin-labeled L43C residues when IM2 TMD was reconstituted in DOPC/DOPS liposomes and native *E. coli* membranes. Generally, the *E. coli* lipid makeup is tuned to accommodate TM helices of the same length as those of IM2. This could explain why the modeled inter-spin label distances based on the NMR structure of IM2 in DOPC/DOPE are much closer to our experimental distances (Figure 4 and Supplementary Figure 5), but large differences were obtained between our DEER distances and these predicted for protein crystals in LPC (Supplementary Figure 5) was observed. The latter could be caused by this special environment-imposed selection of conformations that may be only sparsely populated in the lipids we studied at neutral pH. Therefore, our findings here once again reinforced the role of lipid environment in selecting the protein conformations and possibly tuning its function, which is in general agreement with the current understanding that the lipid environment is not a crowd of passive spectators to the protein but through its properties may tune the conformations and activity of integral membrane proteins.^60,61^

In this work, we also describe a useful protocol for the reconstitution and ESR study of the spin-labeled oligomeric single-pass integral membrane protein IM2 in the *E. coli* native membranes. We find this as an easy and inexpensive alternative to using synthetic lipids and *E. coli* polar extract. Previously, ESR measurements were conducted on endogenously expressed and spin-labeled *E. coli* outer membrane proteins yielding valuable information about the native conformational states of these proteins.^62,63^ But limitations of such studies are substantial, because proteins can be labeled only at accessible cysteine residues in protein regions outside of the cell, and proteins residing in the *E. coli* plasma membrane are almost off-limit to conventional spin labeling practices. The *E. coli* OM is also far enough in its properties from phospholipid bilayer. Therefore, our approach might be useful in investigations of other proteins aiming to test how their structure is affected by the native environment of lipid makeup and membrane proteins.

## AUTHOR CONTRIBUTION

GS: experiment, data analysis and interpretation: PPB: experiment, data analysis and interpretation, writing the manuscript; ERG: conception, design, experiment, data analysis and interpretation, writing the manuscript. All authors read and approved the final version of the manuscript.

## ACKNOWLEDGEMENTS

This study was funded by TTU start-up to ERG. The National Biomedical Resource for Advanced ESR Techniques (ACERT) at Cornell University is funded by an NIH grant 1R24GM146107.

## DATA AVAILABILITY

All experimental data can be provided upon reasonable request to Elka Georgieva and Peter Borbat.

## DECLARATION OF INTERESTS

The authors declare no competing interests.

## Notes

### Competing Interest Statement

The authors have declared no competing interest.

