## Supplementary Information for "A comparative study of influenza A M2 protein conformations in DOPC/DOPS liposomes and in native *E. coli* membranes"

### 1. Reconstitution of spin-labeled IM2 TMD in *E. coli* membrane—further details on sample preparation.

The following analyses and estimations were conducted for the samples of IM2 TMD in native *E. coli* membranes. First, wet *E. coli* cells grown in a 2-liter flask under the conditions described in the **Materials and Methods** (main text) were weighted. Then, the membranes were separated from cell debris, unbroken cells and inclusion bodies by low-speed centrifugation; the membranes found in the supernatant were span down by ultracentrifugation.[1] The wet membranes were collected, weighted, and compared to the weight of *E. coli* wet cells. The weight ratio of wet cells to wet membranes was 5 to 1. Then, the total protein-to-lipid molar ratio (P/L) for IM2 TMD reconstituted in *E. coli* membranes was calculated based on the weight of a single *E. coli* cell of 1 pg or $10^9$ cells per mg of wet cells. On average, one *E. coli* cell contains $2.1\times10^7$ lipid molecules, based on the information from the *E. coli* metabolome database,[2] hence 1 mg of wet cells contains $2.1\times10^{16}$ lipid molecules. Since 1 mg of wet membranes contains 5 times this number, this gives 0.174 µmole phospholipid per 1 mg of wet cells. Based on known concentrations of protein stocks, the estimations of protein-to-lipid ratios for E. coli membrane samples follow. The P/L ratios for all samples are compiled in the Supplementary Table 1. Estimates based on *E. coli* cytoplasmic membrane protein to phospholipid ratios of 0.63 µmol per mg of inner membrane protein[3] give very close estimates for the E. coli phospholipid using respective lipid and protein average weights of 720 and 44,000, and assuming ~35% wet cell hydration, The hydration may vary among different preparations, but is the same for each one, so the P/L ratios are consistent throughout the range.

### 2. Preparation of inner *E. coli* membranes

E. coli inner membranes (IM) were separated from total membranes using a modified EDTA-free sucrose gradient protocol[4] in 26.3 ml screw-cap tubes using Ti 70 rotor at 65 000xg speed for 18 h at 4 °C. The tubes were filled with 38% volume of 73% sucrose solution, 38% of 53% sucrose solution, and homogenized total membranes in 20 mM Tris pH 7.4, 150 mM NaCl and 20% sucrose were loaded on the top to fill the tube volume. After the centrifugation, only the top membrane layer of containing the inner *E. coli* membranes (native lipids and proteins) was collected, washed with buffer without sucrose and span down in the same rotor and tubes at 40k RPM and the membrane pellet was collected.

### 3. Sample compositions and nomenclature

**Supplementary Table1.** Compositions of samples used in this work. Two samples, 1 and 2, of the spin-labeled IM2 TMD were reconstituted in DOPC/DOPS liposomes at final protein-to-lipid molar ratios of 1:480 and 1:230. Three more samples, 3, 4 and 5 of spin-labeled IAM2 TMD were reconstituted in native *E. coli* membranes for protein dilution increasing from 1:1,110 to 1:4,402. Additional preparation of samples 1a to 5a used P/L from 1:416 to 1:10,402. A control sample 6b was made using *E. coli* IM.

| Sample Number | *E. coli* wet membrane weight, (mg), or lipid concentration, (μM), for DOPC/DOPS liposomes | Protein amount added to *E. coli* membranes or final protein concentration in liposome samples | Total P/L ratio |
|---|---|---|---|
| 1 | 11.25 μM | 23.4 μM | 1:480 |
| 2 | 11.25 μM | 47.7 μM | 1:230 |
| 3 | 118 mg | 60 μL × 78 μM | 1:1,110 |
| 4 | 116 mg | 130 μL × 78 μM | 1:1,987 |
| 5 | 112 mg | 230 μL × 78 μM | 1:4,402 |
| 1a | 80 mg | 320 μL × 107 μM | 1:416 |
| 2a | 25 mg | 70 μL × 107 μM | 1:595 |
| 3a | 25 mg | 20 μL × 107 μM | 1:2,080 |
| 4a | 25 mg | 10 μL × 107 μM | 1:4,161 |
| 5a | 25 mg | 4 μL × 107 μM | 1:10,402 |
| 6b(1) | 70.5 mg | 60 μL ×78 μM | 1:2,680 |

(1) *E. coli* inner membranes,

**Supplementary Table2.** Verification of spin label stability to reduction in preparation of E. coli membranes lipid samples.

| Sample number | PE amplitude[a], $A$ (mV) | L/P | DEER modulation depth, Δ | PE dipolar correction, $D=(1-p_A)^{-1}$ | $A\times$LPR$\times D$ $\times10^{-5}$ |
|---|---|---|---|---|---|
| 1a | 500 | 416 | 0.32 | 1.25 | 2.6 |
| 2a | 320 | 595 | 0.27 | 1.21 | 2.3 |
| 3a | 110 | 2,080 | 0.13 | 1.08 | 2.46 |
| 4a | 66 | 4,161 | 0.09 | 1.06 | 2.81 |
| 5a | 25 | 10,402 | 0.05 | 1.03 | 2.7 |

(a) – The concentration could be estimated by using instrumental spin sensitivity factor ~10 mV/µM for nitroxides at current DEER setup. The last column gives spin concentration dependence vs. L/P but is not normalized to yield spin concentrations.

The Supplementary Table 2 estimates spin concentration based on primary echo amplitude $A$ obtained at standard setting. $A$ is scaled up by the dilution factor LPR (Lipid-to-Protein Molar Ratio) and corrected by applying a fudge-factor accounting for PE amplitude loss caused by dephasing of spin-echo due to dipolar coupling. The dephasing is complete within 250 ns pulse separation. An adequate correction used DEER modulation depth Δ, scaled down to $p_A$ corresponding to a softer 32 ns refocusing pulse for A-spins. The main conclusion from the table last column is that the amplitude with is within the rmsd of 7% of the expectation based on estimated spin concentration, which is well within the variations of sample tube volume and protein concentration. This also indicates that if some loss of spin label takes place e.g. to reduction, the effect is insignificant comparing a factor of 6.5 overall change of modulation depth to a maximum factor of 0.21 based on ±1.5 rmsd p-p deviation.

### 4. DEER data for liposomes and E. coli membranes and distance reconstruction

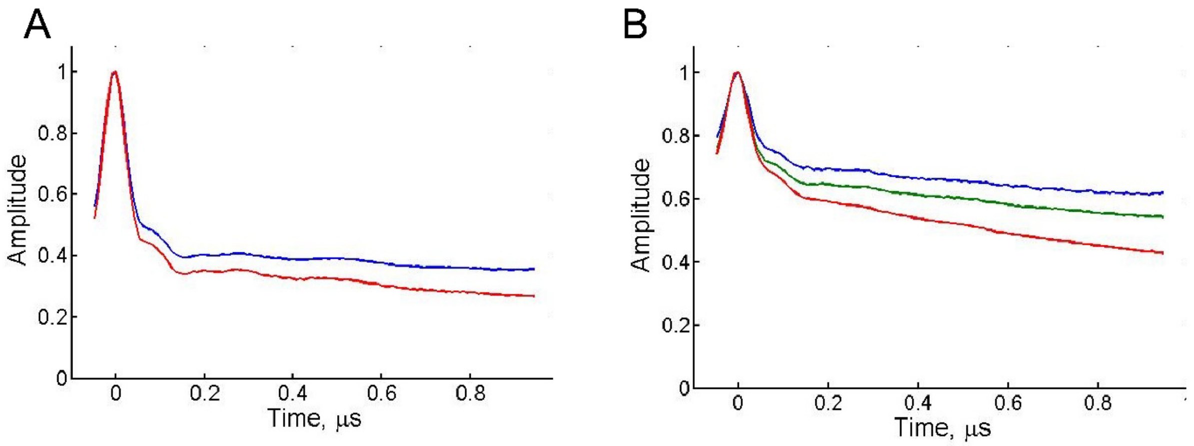

**Supplementary Figure 1.** Raw DEER data for spin-labeled IM2 TMD in (A) DOPC/DOPS liposomes—data for Samples 1 and 2 are in blue and red, respectively; and (B) native *E. coli* membranes—data for Samples 3, 4 and 5 are in blue, green, and red, respectively. Baseline subtraction was conducted as described previously.[5,6]

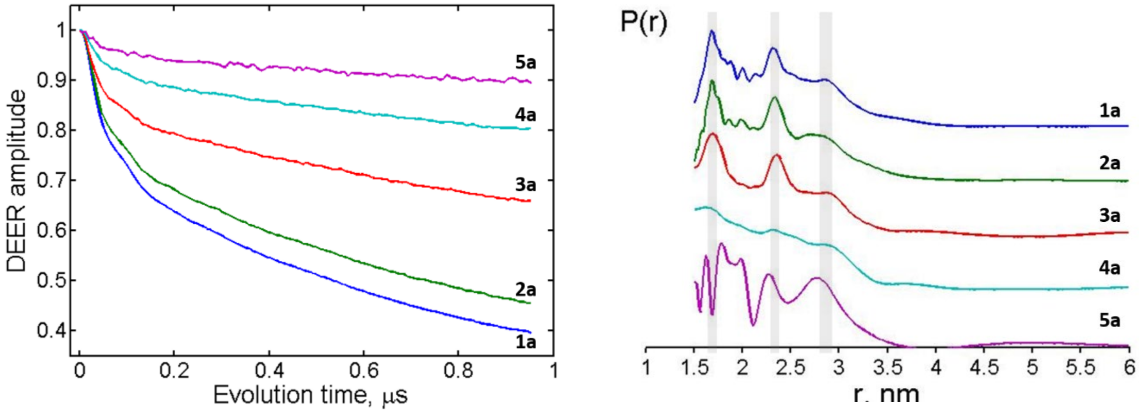

**Supplementary Figure 2.** Raw DEER data for spin-labeled IM2 TMD in native *E. coli* membranes and reconstructed distance distributions—data are for Samples 1a-5a of Supplementary Table 2, as indicated.

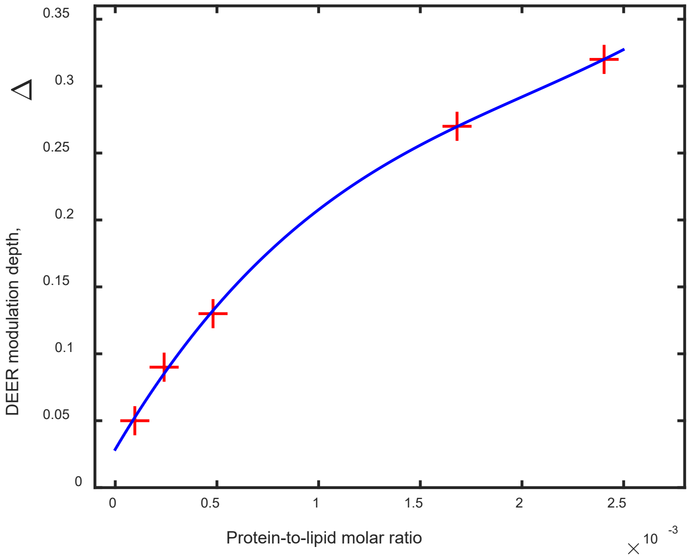

**Supplementary Figure 3**. Binding curve for samples 1a-5a plotted in log scale as a dependence of dipolar amplitude (DEER modulation depth, Δ) vs. P/L molar ratio.

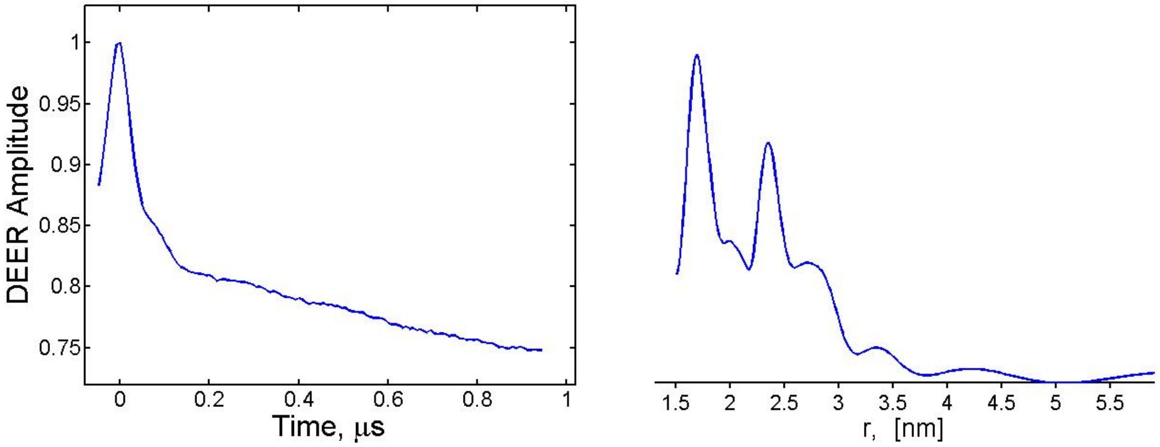

**Supplementary Figure 4.** (Left) Primary DEER data for spin-labeled IM2 TMD in native *E. coli* inner membranes at neutral pH, (Right) Distance distribution produced by Tikhonov regularization using 1st derivative and regularization parameter λ=1.0.

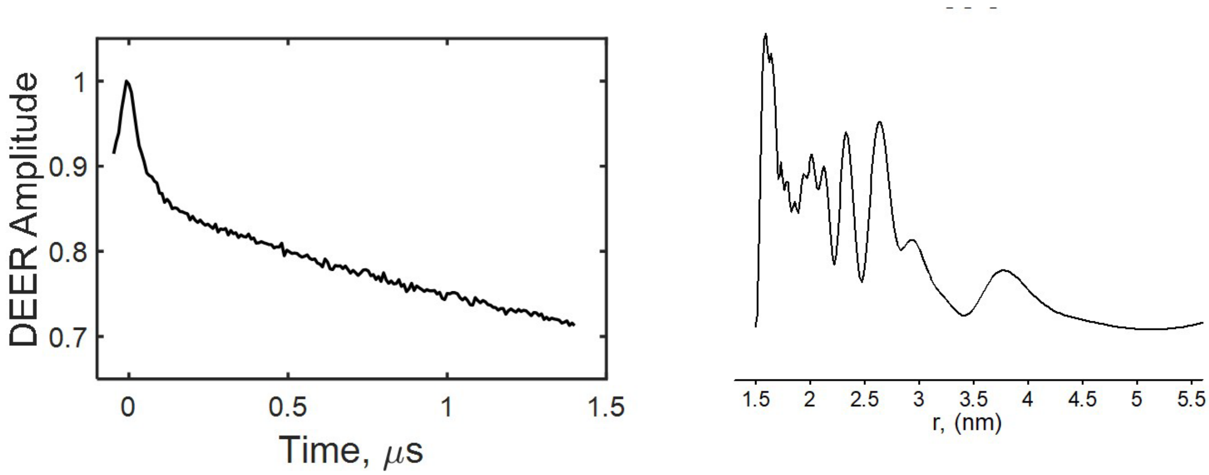

**Supplementary Figure 5.** (Left) Primary DEER data for spin-labeled IM2 TMD in native *E. coli* inner membranes at low pH of 5.4, (Right) Distance distribution produced by Tikhonov regularization using 1$^{st}$ derivative and regularization parameter λ=1.0.

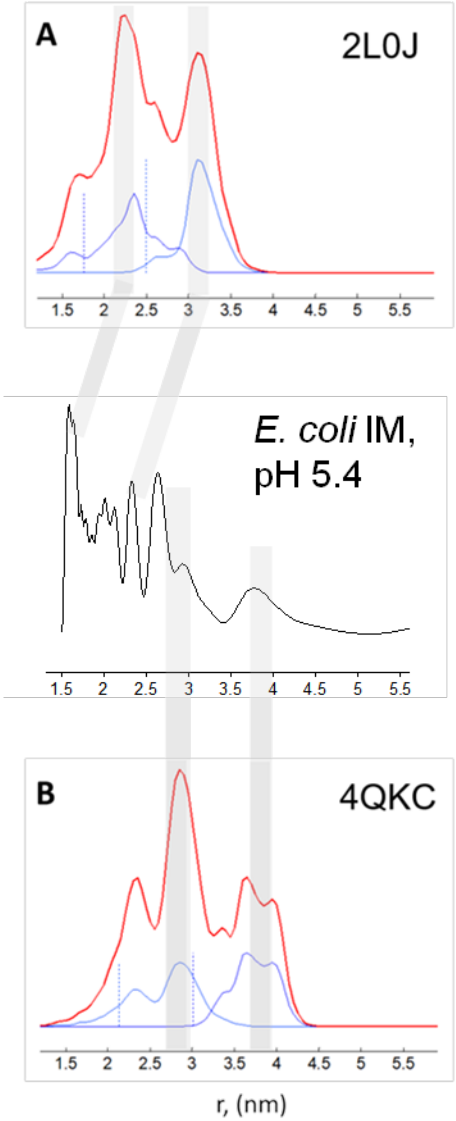

**Supplementary Figure 6.** MMM analysis of distance distributions between spin-labeled residue L43C within IM2 tetramer for (A) NMR (PDB: 2L0J) and (B) X-ray (PDB: 4QKC) structures. The red line corresponds to the total distances and their distributions, whereas the blue lines correspond to the side and diagonal distances in IM2 tetramer. The mid panel compares the distances reconstructed from IM2 in E. coli IM at pH 5.4. The longer distance peaks are comparable in amplitude to shorter distance peak which in turn match peak positions at pH 7.5.

The structures, referred to in the Supplementary Figure 6, were used as templates to generate MTSL rotamer libraries at cryogenic temperature to predict the distances between spin-labels. Based on the set of MTSL rotamers, bimodal distance distributions were obtained with maxima at about 2.2 nm and 3.2 nm for NMR but even longer 2.8 nm and 3.8 nm for X-ray structure. In both cases the distance distribution and distance maxima are significantly different than those which we obtained for IM2 TMD in DOPC/DOPS and native *E. coli* membranes (Figure 3 in the main text) at neutral pH. The rotamer libraries, when built for room temperature, produced broader peaks in the same ranges and are not shown here. The average distances in A are longer than in our experiments. NMR structure would be close, if minor a rotamer producing peak at 1.7 nm would be predominantly populated. The X-ray structure corresponds to a more opened conformation with the distances similar to that in E. coli IM at pH 5.4. Thus, the open structure may reflect a physiological state of IM2.
